# Variance in Estimated Pairwise Genetic Distance Under High versus Low Coverage Sequencing: the Contribution of Linkage Disequilibrium

**DOI:** 10.1101/108928

**Authors:** Max Shpak, Yang Ni, Jie Lu, Peter Müller

**Affiliations:** Sarah Cannon Research Institute, Nashville TN 37203, USA; Center for Systems and Synthetic Biology, University of Texas, Austin TX 78712, USA; Fresh Pond Research Institute, Cambridge MA 02140, USA; Department of Statistics and Data Sciences, University of Texas, Austin TX 78712, USA; Genetics Division, Fisher Scientific, Austin TX 78744, USA; Department of Mathematics, University of Texas, Austin TX 78712, USA

**Keywords:** genetic distance, linkage disequilibrium, coverage, cancer genomics, pooled sampling, next-generation sequencing

## Abstract

The mean pairwise genetic distance among haplotypes is an estimator of the population mutation rate *θ* and a standard measure of variation in a population. With the advent of next-generation sequencing (NGS) methods, this and other population parameters can be estimated under different modes of sampling. One approach is to sequence individual genomes with high coverage, and to calculate genetic distance over all sample pairs. The second approach, typically used for microbial samples or for tumor cells, is sequencing a large number of pooled genomes with very low individual coverage. With low coverage, pairwise genetic distances are calculated across independently sampled sites rather than across individual genomes. In this study, we show that the variance in genetic distance estimates is reduced with low coverage sampling if the mean pairwise linkage disequilibrium weighted by allele frequencies is positive. Practically, this means that if on average the most frequent alleles over pairs of loci are in positive linkage disequilibrium, low coverage sequencing results in improved estimates of *θ*, assuming similar per-site read depths. We show that this result holds under the expected distribution of allele frequencies and linkage disequilibria for an infinite sites model at mutation-drift equilibrium. From simulations, we find that the conditions for reduced variance only fail to hold in cases where variant alleles are few and at very low frequency. These results are applied to haplotype frequencies from a lung cancer tumor to compute the weighted linkage disequilibria and the expected error in estimated genetic distance using high versus low coverage.

## 1 Introduction

One of the defining empirical problems in evolutionary genetics is the measurement and characterization of genetic heterogeneity in natural and experimental populations. The advent of next-generation sequencing (NGS) provides researchers with a tool set for efficiently generating sequence data from large numbers of genotypes and over extensive regions of the genome, including whole-exome and whole-genome sequencing of multiple individuals. This data has the potential to provide the statistical power necessary to make robust inferences of genotype frequencies and their distributions.

High-throughput NGS technology gives researchers choices between different approaches to sampling genotypes from a population. A standard method, most widely used in studies of multicellular organisms, is to sample individuals and sequence their genomes at high coverage, i.e. generating reads containing most or all of the polymorphic sites of interest for each genome. An alternative approach is to sequence from a pooled set of individuals at a read depth much smaller than the number of genomes in the sample, e.g. Futschik and Schlötterer (2010); Anand et al. (2016), leading to a very low average coverage per individual genome. Figure 1 illustrates these two scenarios for a small model population: a sample of *n* individuals sequenced with full coverage, versus low coverage sequencing at read depth *n* from a pooled set of individuals.

**Figure 1.**
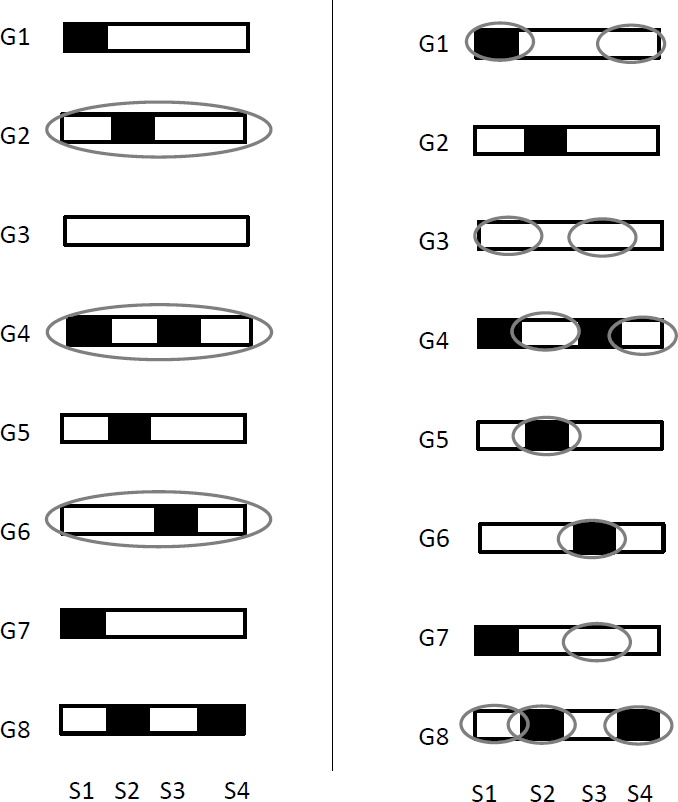
Illustration of high coverage sequencing (HCS) versus low coverage sequencing (LCS). In this example, the population consists of 8 haplotypes G1…G8 characterized by 4 segregating sites *S*1…*S*4. We assume a sampling depth of *n* = 3 and sufficiently many reads to capture all segregating sites. In the left panel, we have a random instance of HCS via the complete sequencing of *G*2, *G*4, *G*5 (gray ovals representing sampling), giving a mean pairwise distance of 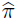 = 2. In the right panel, we have a random instance of LCS, such that G1, G3, G8 are sequenced at S1, G4,G5 and G8 at S2, etc, giving a mean genetic distance 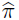 = 8*/*3. Note that *E*(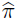) is the same under both modes of sampling, the differences are due to *var*(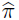).

Sequencing at low coverage is typically used in population genetic studies of microbial assemblages and in cancer genomic studies where genetically heterogeneous assemblages of cancer cells are sampled from a single tumor. However, through single-celled sequencing techniques (Navin, 2015; Gawad et al., 2016), individual sampling with high coverage is also possible for these model systems. Similarly, while individual sampling has been standard in population genetic studies of most multicellular organisms, NGS has made pooled sampling with low coverage sequencing inexpensive and practical in studies of animal and plant populations. For example, several recent analyses of genetic variation in *Drosophila* populations (Schlötterer et al., 2014) used low coverage pooled sequencing, drawing reads from a very large pool of macerated flies rather than sequencing fly genomes individually with high coverage.

Sequencing *n* individuals with full coverage is not statistically equivalent to sequencing at read depth *n* from a large pool of individuals. High and low coverage result in different estimation error for population parameters. These include the population mutation rate *θ* = 4*Nu* (where *N* is the population size and *u* the genomic mutation rate, with *θ* = 2*Nu* for haploid genomes), which is estimated either from the number of segregating sites (Watterson, 1975) or from the average heterozygosity across sites (Tajima, 1989). Estimates of *θ* are the basis for a number of statistical tests that distinguish the effects of natural selection and population dynamics from neutral evolution at constant population size. These include the Tajima‘s D test (Tajima, 1989), which compares *θ* estimates from the number of segregating sites to those derived from average heterozygosity. Consequently, getting a handle on the variance in estimates of *θ* and for neutrality test statistics generally is of broad interest and importance in evolutionary genetics (Nielsen, 2001). Several studies have analyzed the contributions of pooling, read depth, and coverage to bias and variance in *θ* estimates, e.g. Pluzhnikov and Donnelly (1996); Lynch (2008). For example, given a con-5 stant read depth, pooling improves the accuracy in estimated *θ* due to effectively larger sample size (Futschik and Schlötterer, 2010; Ferretti et al., 2013), while Korneliussen et al. (2013) have shown that low read depth can lead to estimation bias in the Tajima D test statistics.

Considering the effects of coverage on parameter estimation, if the number of genomes sampled is held constant, lower coverage leads to smaller sample size, and consequently greater error. However, Ferretti et al. (2014) have shown that as long as the reduction in coverage is compensated by the number of genomes represented in a sample, low coverage sequencing reduces the error in estimates of *θ* and the Tajima D statistic. Specifically, if we estimate allele frequencies and *θ* from *n* sequences with complete coverage, as opposed to a much larger number of sequences at very low per-genome coverage (so that on average each site is represented by *n* samples, often from different individuals per site, as shown in panel 2 of Fig. 1), low coverage sequencing reduces the error in estimated *θ*. Ferretti et al‘s results are explained by the fact that with low coverage sequencing, variant alleles from different segregating sites tend to be sampled from different individuals, corresponding to an effective increase in the number of independent genealogies from which variant allele samples are drawn for each locus. Consequently, their results imply that the degree to which estimates of *θ* improve with low coverage, large individual sample sequencing are expected to increase with the strength and direction of linkage disequilibria among polymorphic sites.

In this study, we will consider limiting cases of high and low coverage sequencing to investigate the contribution of linkage disequilbria to estimates of *θ*. High coverage sequencing (HCS) is represented by complete coverage of all polymorphic sites from *n* different genomes, as would typically be the case for individual sampling (including single-cell sequencing). Low 6 coverage sequencing (LCS) is represented by a case where a very large sample of genomes is pooled and sequenced at a read depth *n* for each site, so that allelic variants at different sites are almost always drawn from different genomes. We will compute variances in the Tajima estimator *E*(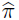) = 4*Nu* = *θ*_*π*_, which is calculated from the mean pairwise genetic distance in a sample of *n* genotypes:

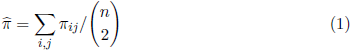

where 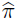_i,j_ is the Hamming distance for the haplotype pair *i, j* summed over all polymorphic sites.

We hypothesize that under most conditions, the variance in 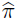 estimated using HCS increases with greater linkage disequilibrium across polymorphic sites, i.e. strong linkage disequilibria inflate the estimation error across sites in a haplotype by reducing the number of independent genealogical sample paths. We will investigate this hypothesis analytically, and will additionally validate our results using individual-based simulations. We will also apply these results to NGS data by analyzing allele and haplotype frequencies from cancer cell genomes.

## 2 The Sampling Models

Consider a population of *N* organisms with mutations distributed over *S* segregating sites. We wish to estimate the mean genetic distance population and its sample variance *var*(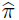 for the 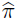) under the high and low coverage modes of sequencing. For HCS, we draw *n ≪ N* individual organisms (or cells) from the population and sequence their entire genomes, exomes, or any regions containing the polymorphic sites of interest.

For an idealized model of LCS, we assume a mean coverage depth *n ≪ M*, where *M* is the number of genotypes contributing to the pooled sample (*M* may be *≪ N* or of the same order). If reads are short, the majority will contain at most a single polymorphic site. Together, these conditions lead to each polymorphic site being sampled independently of other polymorphic sites with respect to the genome of origin (note that in the second panel of Figure 1, multiple sites are sampled from the same genome simply because there are very few genomes to draw this random sample from). When computing sample genetic distance, extreme HCS sums over the Hamming distances of all haplotype pairs, while extreme LCS results in summing over all pairs for each segregating site sampled from a different genome.

We assume an infinite sites model (Kimura, 1969; Tajima, 1996) so that there are only two alleles per segregating site. This allows an unambiguous binary classification of alleles, with mutations as ancestral "wildtype" vs. "reference" genotype. This also allows us to specify the direction (sign) of linkage disequilibria. For cancer cells, the reference corresponds to the normal germline genotype, with somatic mutations defining the variant genotypes of the clonal lineages. Our methods and results also apply to multiallelic states provided that some allele, usually the most common, is designated as a reference and all other alleles are aggregated to create a biallelic state.

**Definitions**. Throughout the paper, we use the following definitions and terminology as a formal way of defining Eqn. (1) for high and low coverage sequencing:

*Variables*: Let *z* denote a genotype, at either a single locus *s* or across multiple loci. We define the frequency distribution of *z* over samples *i* as *z*_*i*_ ~ *p*(*z*), which are iid among *i* = 1…*n*. We use *z*_*is*_ to denote site *s* in haplotype *i* (for HCS). We write *z*_*i_s_,s*_ to denote sample *i* at site *s* when sites are sampled by pooling at low coverage (LCS) when we wish to highlight the fact that *z*_*is*_ and *z*_*ir*_ are read from distinct haplotypes.

*Pairs*: For both HCS and LCS, the estimators of 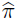 include an average *Σ_i<j_ ϕ_ij_/n*(*n* − 1) of some function *ϕ*_*ij*_ = *ϕ*(*x_i_, x_j_*) of pairs of i.i.d. random variables *x*_*i*_, *i* = 1, …, *n*. In the case of LCS *x*_*i*_ = *z*_*is*_ and *ϕ*(*x_i_, x_j_*) = *f*_*ijs*_ with *f*_*ijs*_ = *I*(*z_is_* ≠ *z*_*js*_)(and an additional sum appears over *s*, outside the average). For HCS the random variables are *x*_*i*_ = ***z****_i_* and *ϕ*(*x_i_, x_j_*) = *g*_*ij*_ = *Σ_s_ f_ijs_*. Importantly, while the *x*_*i*_ are independent, pairs (*x_i_, x_j_*) and (*x_i_, x_k_*) that share a common element are not, and thus the same for *ϕ*_*ij*_.

*Moments of ϕ*_*ij*_: We define *E*(*ϕ*_*ij*_) = *µ*, *var*(*ϕ*_*ij*_) = *σ*^2^. We also define an expectation for the product *E*(*ϕ_ij_, ϕ_jk_*) =*κ* for pairs of pairs with a shared element.

*Pairs of pairs*: Let *P* denote the set of all ordered pairs of pairs, with *P*_3_ *⊂ P* defining the subset of ordered pairs of pairs with a single shared element,

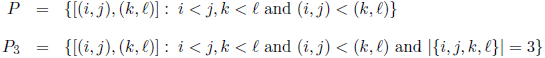

*Numbers of pairs*: The number of ordered pairs, and the number of ordered pairs of pairs with a shared element are, respectively *N*_2_ = *n*(*n* − 1)*/*2 and *N*_3_ = *n*(*n* − 1)(*n* − 2)*/*2. The value of *N*_3_ follows from there being *n*(*n* − 1)(*n* − 2)*/*6 ways to select a triplet *i, j, k*, and three ways to select a shared element from this triplet. In Appendix A1, we discuss the properties of ordered pairs of pairs, including the derivation of the following relation which we will use below to compute *var*(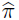) under low and high coverage sequencing,

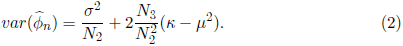

where 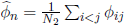 is a sample estimate of *E*(*ϕ*_*ij*_) = *µ*. We will use this result twice, once for LCS with *ϕ*_*ij*_ = *f*_*ijs*_, and once for HCS with *ϕ*_*ij*_ = *g*_*ij*_.

### 2.1 Case 1: Low Coverage Sequencing (LCS)

For LCS, we use the indicator function at a single site *s*, *f*_*ij,s*_ = *I*(*z_i_s_,s_* ≠ *z*_*js,s*_), where *z_i_s_,s_ ~ Bern*(*p*_*s*_), i.e. 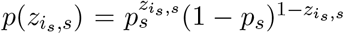 for *z_i_s_,s_ ∈ {*0, 1*}* such that

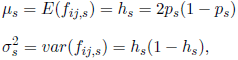

where *h*_*s*_ is also known as the heterozygosity at locus *s*. Under LCS we define the estimator for 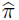 for Eqn. (1) as:

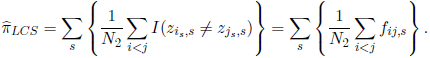

To evaluate the variance of 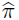_LCS_, first note that the expectation of products *f_ij,s_, f_jk,s_* for ordered pairs is:

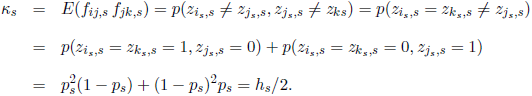

From the assumption of statistical independence among sites *s* located on different reads with pooling, it follows (Appendix A1) that for a sample of size *n*,

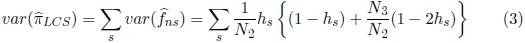

Approximate statistical independence across sites requires that the number of possible samples of size *n* is much larger than the number of segregating sites (i.e. *N ˛ n* so that 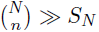).

### 2.2 Case 2: High Coverage Sequencing (HCS)

Computing pairwise differences among samples of individuals genomes under HCS involves calculating moments of sums rather than sums of moments. For **z***_i_ ~ p*(**z**) sampled independently under HCS, applying 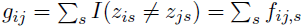, we define

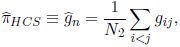

and derive its variance below. For a sample of individual haplotypes *i* = 1…*n*, consider *z_is_ ~ Bern*(*p*_*s*_) as before, but with correlated *z_is_, z_ir_* due to linkage disequilibrium (LD) between sites. We define the LD *D*_*rs*_ for (arbitrarily labeled) alleles *R, r* and *S, s* at the two sites as follows. Letting *q_s_, q_r_* = 1 *− p_s_,* 1 *− p*_*r*_ (Lewontin and Kojima, 1960),

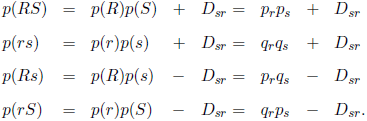

As with LCS, we have, for *h*_*s*_ = 2*p_s_q_s_*,

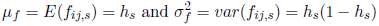

The probability of different identity among sites *s, r* in a sample pair *i, j* is *p*(*f_ijs_f_ijr_* = 1) = *p*(*RS, rs*) + *p*(*rs, RS*) + *p*(*Rs, rS*) + *p*(*rS, Rs*), where *p*(*RS, rs*) = *p*(*z*_*i,sr*_ = *RS, z*_*j,sr*_ = *rs*) etc. Therefore

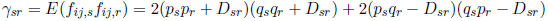

and similarly, considering triplet samples with shared element *j* paired with *i* and *k*, the probability of different identity between *i* and *j* at site *r* and *j* and *k* at site *s* is *p*(*f_ijs_f_jkr_* = 1) = *p*(*R, rS, s*) + *p*(*R, rs, S*) + *p*(*r, RS, s*) + *p*(*r, Rs, S*), where *p*(*R, rS, s*) = *p*(*z*_*ir*_ = *R, z*_*jr*_ = *r, z*_*js*_ = *S, z*_*ks*_ = *s*) etc. Using these terms, we compute the expectation:

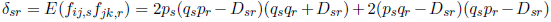

At linkage equilibrium (*D*_*sr*_ = 0 for all *s, r*), high coverage sequencing of *n* individuals and low coverage sequencing of pooled individuals at read depth *n* are statistically equivalent, i.e. both equations simplify to *γ*_*sr*_ =*δ*_*sr*_ = 4*p_s_q_s_p_r_q_r_*.

The mean and sample variance terms for the expected pairwise distances are, respectively,

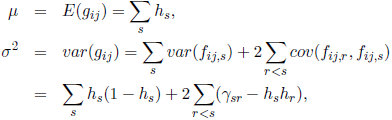

while the covariance 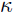 for the ordered pair of pairs with a shared *j* element is:

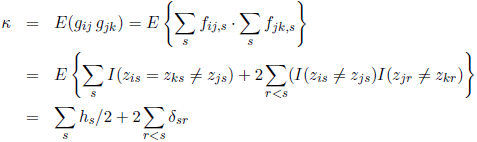

By incorporating 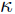, we construct the sample estimate and variances for *g*_*ij*_. Because we are now averaging over haplotypes **z***_i_* (rather than independent counts for each site), 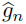 is itself an average across pairs, like 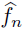 in the LCS case. Applying Eqn. (2), we find that

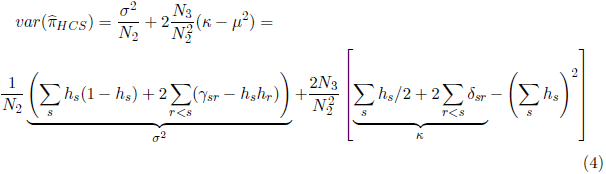

### 2.3 Difference and Independence

Using the results in Eqns. (3) and (4), we derive the difference between the sample variances in pairwise differences under HCS vs. LCS as

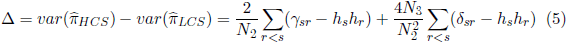

By collecting terms, we can rewrite the above as

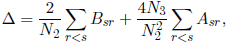

where

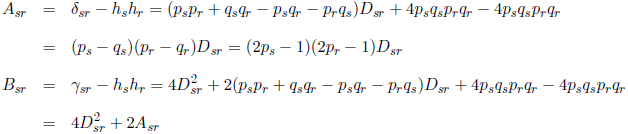

For notational convenience, we define:

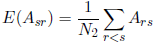

Without linkage disequilibria among pairs (*D*_*sr*_ = 0 and therefore *A_sr_, B_sr_* = 0 for all *s, r* pairs), *γ*_*sr*_ =*δ*_*sr*_ = *h_s_h_r_* and Δ = 0, i.e. the sample variances under HCS and LCS are equal. Because *B_sr_ ≥ A_sr_* for *A_sr_ >* 0, *E*(*A*_*sr*_) *>* 0 is a sufficient condition for Δ > 0. This condition is satisfied when the sum of weighted linkage disequilibria *A*_*sr*_ is positive, i.e.

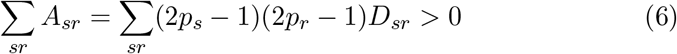

We remark that *E*(*A*_*sr*_) *>* 0 is a sufficient but not necessary condition for Δ *>* 0. The variance in mean pairwise distance can be reduced with pooled LCS even for *E*(*A*_*sr*_) *<* 0, because negative *A*_*sr*_ may be offset by the positive contributions of *D*^2^*_sr_* to the *B*_*sr*_ term when pairwise LD values in the population are sufficiently high. However, with large sample sizes, the *A*_*sr*_ term dominates because it scales as ~ 1*/n* while the *B*_*sr*_ term scales as ~ 1*/n*^2^; consequently, the sign of *E*(*A*_*sr*_) generally predicts that of Δ.

*E*(*A*_*sr*_) *>* 0 requires that most "major" alleles (those with *p_s_, p_r_ >* 0.5) at different loci are in positive LD, while major and minor allele pairs at different loci (*p_s_ >* 0.5, *p_r_ <* 0.5 or vice-versa) are in negative LD. The weighted LD *A*_*sr*_ provides a measure of the extent to which major alleles are in positive LD, regardless of whether the more common allele is a reference/wildtype or variant/mutant at a particular site. These results predict that when the mean weighted LD is positive, the sample variance (error) in estimated pairwise genetic distance will be reduced by LCS.

#### 2.3.1 Possible Caveats: Random Read Depth and Fractional Coverage

Our results are based on a comparison of two extreme-case scenarios: full coverage sequencing of *n* sampled genomes versus low coverage sequencing of *M* sampled genomes at a fixed read depth *n ≪ M*, which is an idealization, because in reality the actual read depth under NGS varies across sites. ***Random Read Depth***. We first consider the effect of having the read depth as Poisson random variable rather than a constant. In Appendix A2, we show that if the read depth *n_s_ ~ P oiss*(*n*) (for mean read depth *n*), then 15 the variance of the estimated pairwise genetic distance under LCS is:

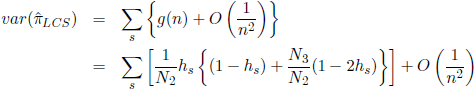

This result follows because a Poisson random variable with mean *n* has variance = *n* (indeed, read depth variance across sites will be typically of *~ n* for most other plausible sampling distributions with a mean read depth *n*), the contributions of the variance in read depth to the variance of the estimate are *~ O*(1*/n*^2^), meaning that the *var*(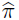*_LCS_*) is essentially the same as Eqn. (3) and *A*_*sr*_, so that Δ remain essentially unchanged. Appendix A2 also presents numerical examples confirming these results.

***Fractional Coverage***. Our derivations for LCS assumed that the coverage in LCS was sufficiently low that effectively only a single locus is sampled per individual, which is an idealization for very short reads and for very large, well mixed pooled sample size *M* relative to the read depth. With greater coverage, LCS results in the sampling of several polymorphic loci from within one genome, albeit far fewer than complete coverage sequencing of individual genomes. We show that the variances in 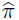 with reduced coverage will be between the limiting LCS case and HCS variances based on a continuity argument. To see the effect of a mode of sequencing where a fraction *ρ* of each genome is covered, consider the covariance term *γ*_*sr*_ used in the derivation of Eqn. (4), *γ*_*sr*_ = *E*(*f_ij,s_f_ij,r_*). Let *ξ*_*i*_ = *I*(*z*_*is*_ and *z*_*ir*_ are phased) denote an indicator for recording phased alleles identified from a single haplotype (such as through the use of long reads).

Different (*ξ_i_, ξ_j_*) result in different expressions for *γ*. If both are on the same haplotype, *ξ*_*i*_ = *ξ*_*j*_ = 1, we get the *γ*_*rs*_ in the HCS limit (see derivation of Eqn. (4)). If both are independent, *ξ*_*i*_ = *ξ*_*j*_ = 0, we get 2*p*_*s*_(1 *− p*_*s*_) *·* 2*p*_*r*_(1 *− p*_*r*_) = *h_s_h_r_* in the LCS limit. We apply the law of total probability to evaluate

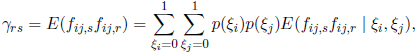

with *p*(*ξ*_*i*_ = 1) = *p*(*ξ*_*j*_ = 1) = *ρ*. The same argument from total probability (applied over cases where *i, j, k* have *s, r* sites on various combinations of shared vs. different reads) applies to continuity of *γ*_*rs*_.

Let 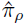 denote the estimator under a sampling scheme with fractional coverage. It follows from the continuity of *δ, γ* and from the fact that the marginal expectations *E*(*f*_*ij*_) are the same under HCS and LCS that the resulting var(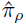) is a continuous function of *ρ* with 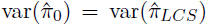 and 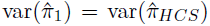. This result is qualitatively consistent with the relationship between the variance in *θ* and the fraction of missing sequence data in Ferretti et al. (2014).

## 3 Dependence on Allele Frequencies and Linkage Disequilibria

Low coverage sequencing reduces the error in estimates of 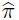 when Δ *>* 0, which holds if the expected weighted linkage disequilibria defined by Eqn.(6) are positive. Ferretti et al. (2014) observed a decrease in variance under low coverage for populations at mutation-drift equilibrium - their result suggests that *E*(*A*_*sr*_) should be positive for equilibrium distributions of allele frequencies in an infinite sites model. Below, we show that this condition holds provided there is a sufficiently high probability density of high frequency mutations, i.e. sufficiently many haplotypes with multiple mutations.

As noted above, *E*(*A*_*sr*_) *>* 0 requires positive linkage disequilibria among pairs of alleles that are rare (*p <* 0.5) and among pairs of alleles that are common (*p >* 0.5), and by symmetry, negative LD among most pairs of major/minor alleles across loci. For a population of clonal, non-recombinant genomes, we can determine the distribution of LD given a distribution of allele frequencies. In an infinite sites model without recombination, letting *p_s_, p_r_* be the frequencies of variant alleles *S, R* at loci *s, r*, where *p_s_ < p_r_*, it follows that all occurrences of *S* must be in *S, R* haplotypes (conversely, if *S* co-occurs with *r*, there can be no *S, R* haplotypes). Therefore, if *p*_*s*_ + *p_r_ >* 1 and *p_s_ < p_r_*, then *p*(*S, R*) = *p*_*s*_ and the LD between loci *s, r D*_*sr*_ = *p*_*s*_(1*−p*_*r*_). When *p*_*s*_ +*p_r_ <* 1, we can compute the expected LD by counting the number of cases that *S* can co-occur with the *R* allele versus the *r* wildtype, since the constraint of *S* either always or never co-occurring with *R* when *p_s_ < p_r_*.

### 3.1 Computing Linkage Disequilibria and *E*(*A*_*sr*_)

Let *N* be the total number of haplotypes and let *s, r* denote any two loci. We observe variant allele frequencies 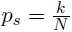 and 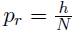 at loci *s* and *r* for *k, h* = 1, …, *N*. Without loss of generality, we assume *k ≤ h* (corresponding to *p_s_ < p_r_*). Let *C*_*sr*_ denote the event that mutations *S* and *R* co-occur and *S* and *r* don‘t co-occur, while 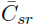 denotes the event that *S* and *R* don‘t co-occur (i.e. *S* co-occurs with the "wildtype" allele at *r*). Let 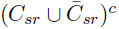 denote the complement of 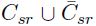 such that *S* and *R* co-occur on some haplotype and *S* and *r* co-occur on some other haplotype. Because there is no recombination and no multiple mutations per site, 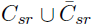 and 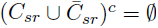. We compute the probability of co-occurrence 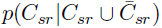:

(1) **N < h + k**. Trivially, 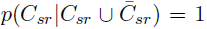 and 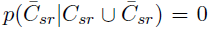, i.e. *p_r_ + p_s_ > 1 → p(C_sr_) = p_s_*.

(2) **N ≥ h + k**. We first derive *p(C_sr_)* and 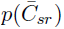 and then 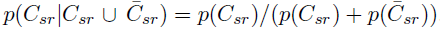.

Counting the possible combinations, we find

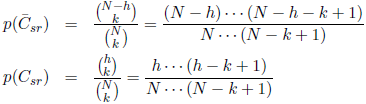

The numerators in the first and second equations give the number of ways in which *k S* alleles can co-occur with *r* or with *R*, respectively, given 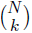 possible positions for the *S* alleles. Therefore,

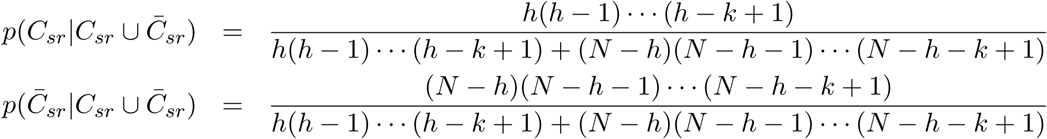

Given *C*_*sr*_, *p*(*SR*) = *p*_*s*_ = *k/N* and hence 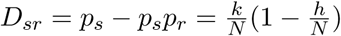.

Given 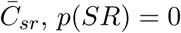 and hence 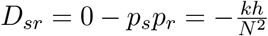.

Therefore, for 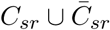,

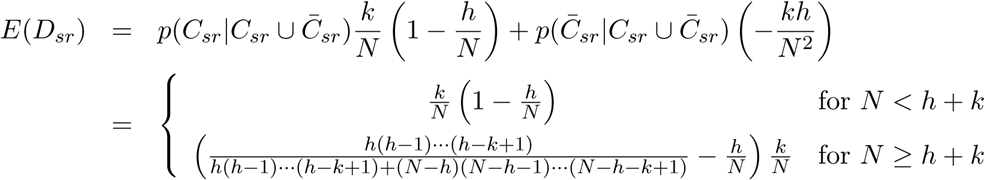

As expected, in the limiting cases of a new mutation (*k* = 1), *D*_*sr*_ = 0, while for *k* + *h > N*, *E*(*D*_*sr*_) = *p*_*s*_(1 *− p*_*r*_).

To compute *E*(*A*_*sr*_), we evaluate the weighted linkage disequilibria over the distribution of *η*_*k*_, *η*_*h*_, the number of sites at which there are exactly *k, h* copies of variant alleles. The expected weighted LD***η*** = (*η*_1_, …, *η*_*N*_) is

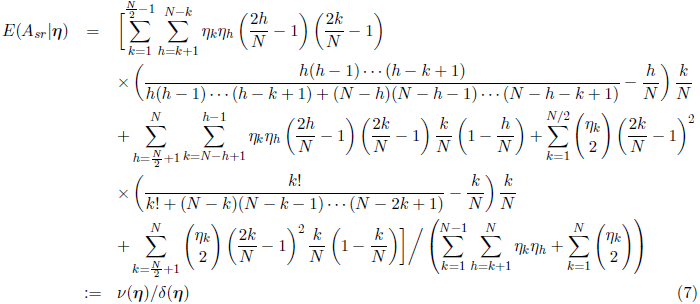

with the convention that 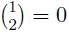, so that the marginal expectation of *A*_*sr*_ is given by

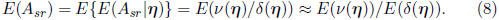

This approximation holds based on a Taylor expansion of the ratio of two random variables, and the fact that *v(**η**)* is much smaller than *δ(**η**)* due to the typically small values of *A*_*sr*_ for any pair *s, r*. The ratio of expectations is:

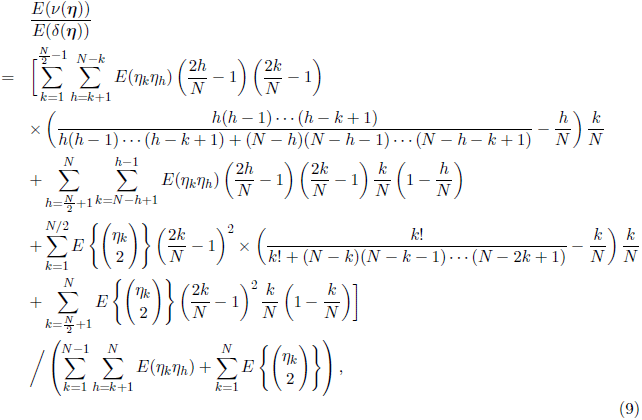

The expectations 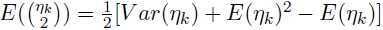, while 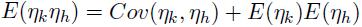. Therefore, we can approximate the expectation of *E*(*A*_*sr*_) (and at least obtain the correct sign from the numerator) to determine if Δ is positive when the expectations and second moments of an allele frequency distribution are known. The same approach can be used to compute the expectated linkage disequilibria over all *η*_*k*_, *E*[*E*(*D_sr_|****γ***)], by substitution *D*_*sr*_ for *A*_*sr*_ (i.e. no factor of (2*p*_*s*_ − 1)(2*p*_*r*_ − 1)) in the equations above.

To calculate *E*(*A*_*sr*_) for an infinite sites model at mutation-drift equilibrium, we use the expectations and variances of *η*_*k*_ (Fu, 1995):

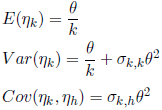

where the coefficients *σ*_*k,k*_ and *σ*_*k,h*_ are functions of harmonic series sums *a*_*n*_ and *B*_*n*_, i.e.

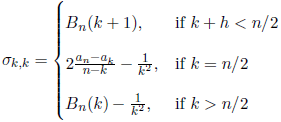

while the covariance coefficients are (for *h > k*)

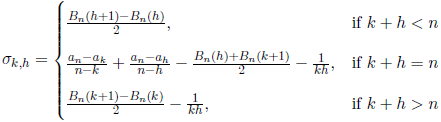

where

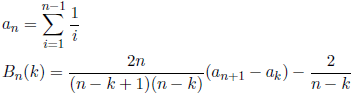

Ferretti et al. (2014) used the expectations and covariances of *η*_*k*_ to derive sample variances for *θ* and the Tajima D test statistic, showing that these variances were reduced with LCS for equilibrium allele frequency distributions under an infinite sites model. Using the expectations for (weighted) linkage disequilibria, we show that these results follow as consequences of the distributions of pairwise LD, and compare these predictions to simulation results for representative parameters. We remark that numerical estimates of *E*(*A*_*sr*_) computed from Eqn. (9) will be only approximate for several reasons: first, because the expectations of ratios do not exactly equal ratios of expectations (Eqn. (8)), and second, because the expectation and variance of *η*_*k*_ are derived for sampling distributions where *n ≪ N* (Watterson, 1975; Fu, 1995). However, they usually provide a sufficiently good approximation (Wakeley and Takahashi, 2003) as *n → N*, so that our estimates of *E*(*A*_*sr*_) should be of the correct sign and magnitude.

## 4 Comparison to individual-based simulations

To simulate Fisher-Wright genetic drift in an infinite sites model, we initialized a population of *N* haploid genotypes at *K* = 10^8^ sites with reference genotypes. In every generation, *N* individuals were sampled with replacement from the existing pool, with each individual sampled producing a single progeny. The number of mutations *m* for each offspring is *m ~ P oiss*(*Ku*), with the mutations randomly distributed among the *K* sites. This process was iterated over *T* generations; in order to approximate a near-equilibrium distribution of allele frequencies, we ran the simulations for *T ~* 4*N* to assure a coalescent among all lineages in each sample path genealogy. Simulations were also run for a range of values *T < N* to generate non-equilibrium distributions of allele frequencies and pairwise LD. For each combination of parameters, the simulation cycle was replicated 100 times.

To simulate full coverage sequencing of individuals, *n* haplotypes were randomly selected without replacement from the model population. The Hamming distances were calculated for all pairs in a sample, variant allele frequencies and linkage disequilibria were calculated over all individuals and all pairs in the model population. Sequencing of pooled samples at very low coverage was simulated by selecting *n* alleles without replacement at every segregating site and summing pairwise distances over all sites (corresponding to sampling with replacement with respect to genomes, but without replacement with respect to each locus). Δ was estimated from the data as the difference in the sample variances between the HCS and LCS pairwise distances. In every replicate run, *A*_*sr*_ was calculated from the mutation frequencies *p_s_, p_r_* and *D*_*sr*_ using Eqn. (6). All simulations were implemented using Python 2.7.3, the code is available from the corresponding author upon request.

The simulation results for population sizes *N* = 200, 500, a sample size of *n* = 20 and a range of generation times *T* are shown in Tables 1 and 2. The first table shows the estimated parameter values from which Δ is calculated, including the number of polymorphic sites *S*_*N*_ in the population (as opposed to the sample number of segregating sites *S*_*n*_), the population mean allele frequency across polymorphic sites, the sample mean pairwise genetic distances under HCS and LCS (for *n* = 20), and their sample variances over 100 replicates.

**Table 1a-b.**
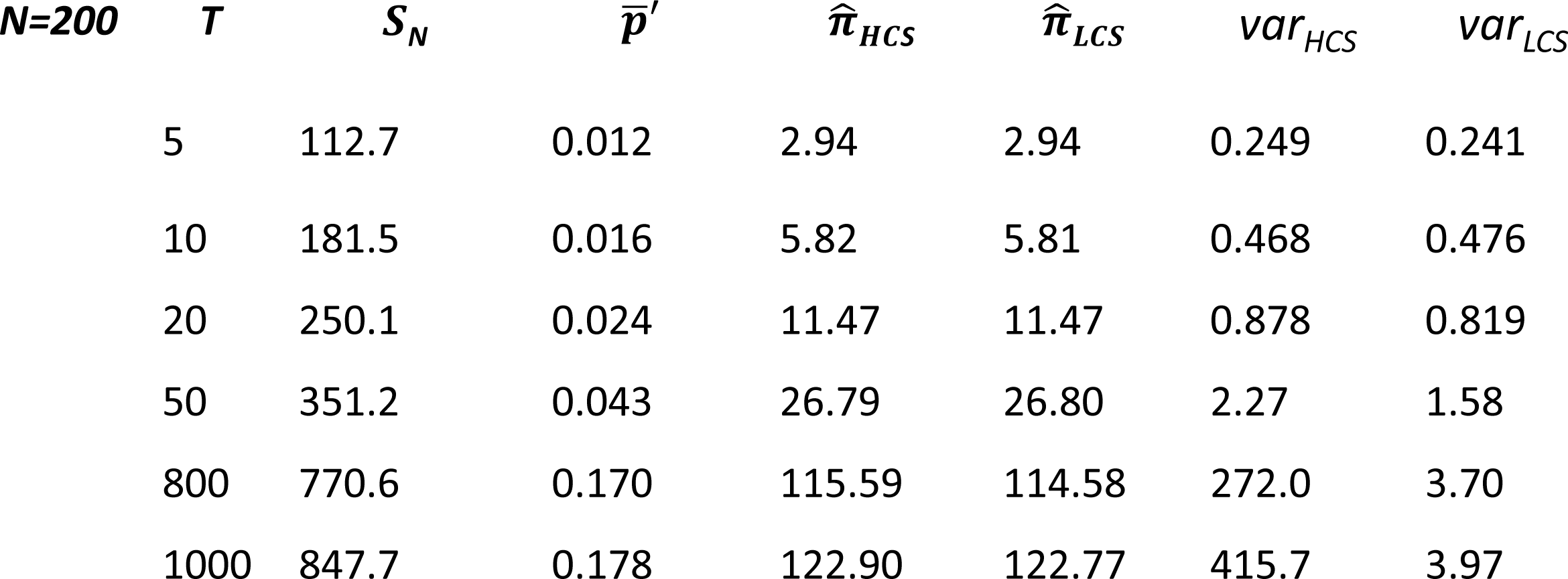
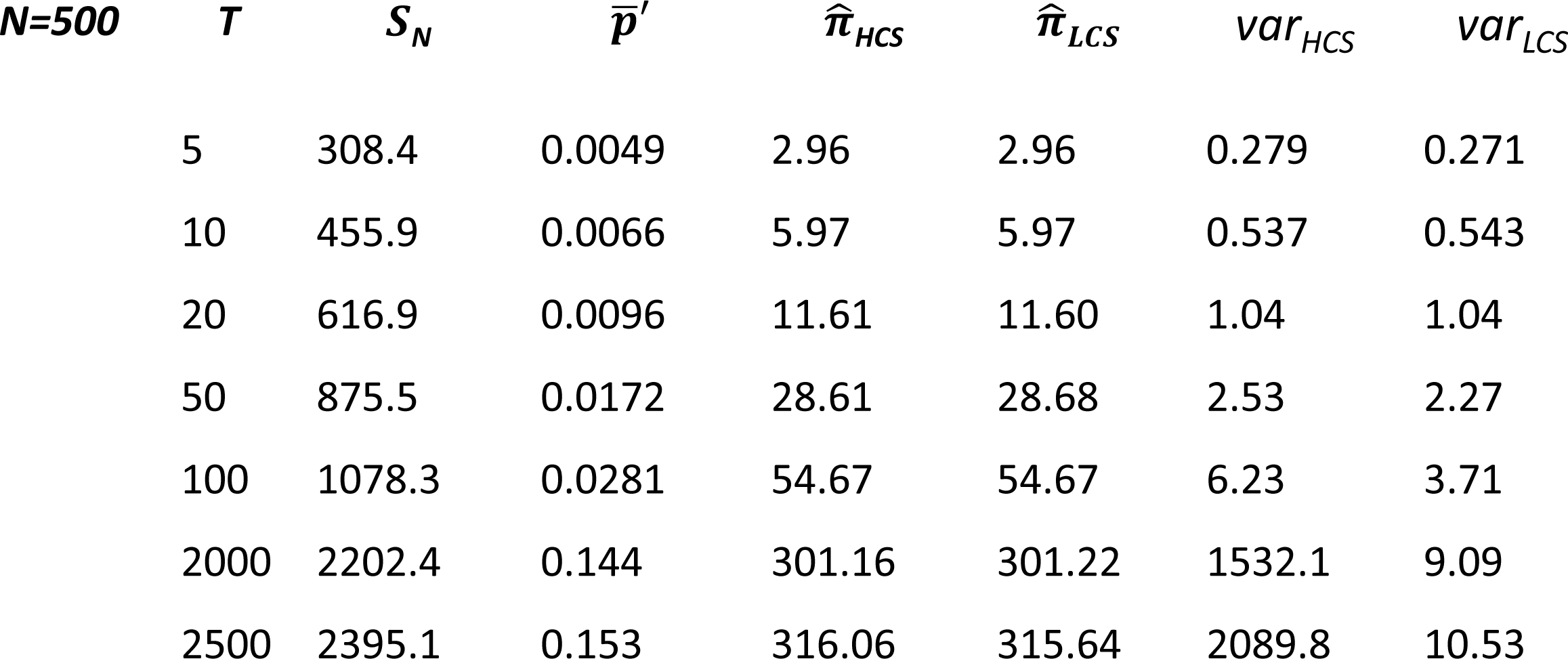
A summary of population parameters for a simulated Fisher-Wright model of genetic drift with infinite sites. The table shows a comparison of Δ*_P_* values predicted from Eqn. (5) with simulation the values Δ*_S_* for *N* = 200 (Table 1a) and *N* = 500 (Table 1b) and sample size/coverage depth *n* = 20 for a range of time intervals, including values near *T* = 4*N* close to equilibrium. The standard error of Δ*_S_* is also shown, estimated Δ*P* is within *<* 2(*SE*) units from Δ*_S_* even for small *T* and few mutations. Mean population pairwise LD values (not shown) are all essentially zero for all simulations, while the magnitudes of *A*_*sr*_ increase with *T* as predicted. *p* is the mean variant allele frequency across all segregating sites.

**Table 2a-b.**
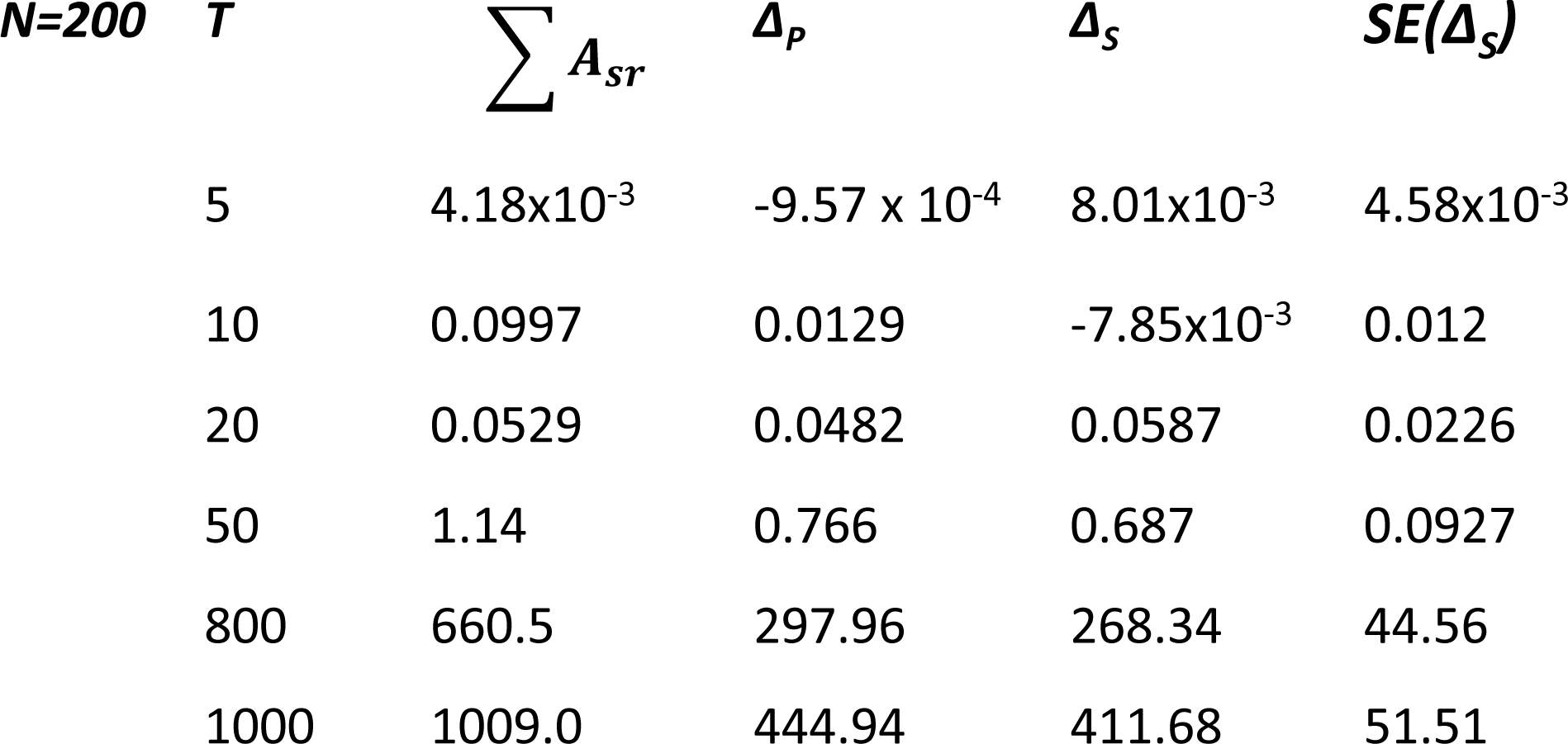
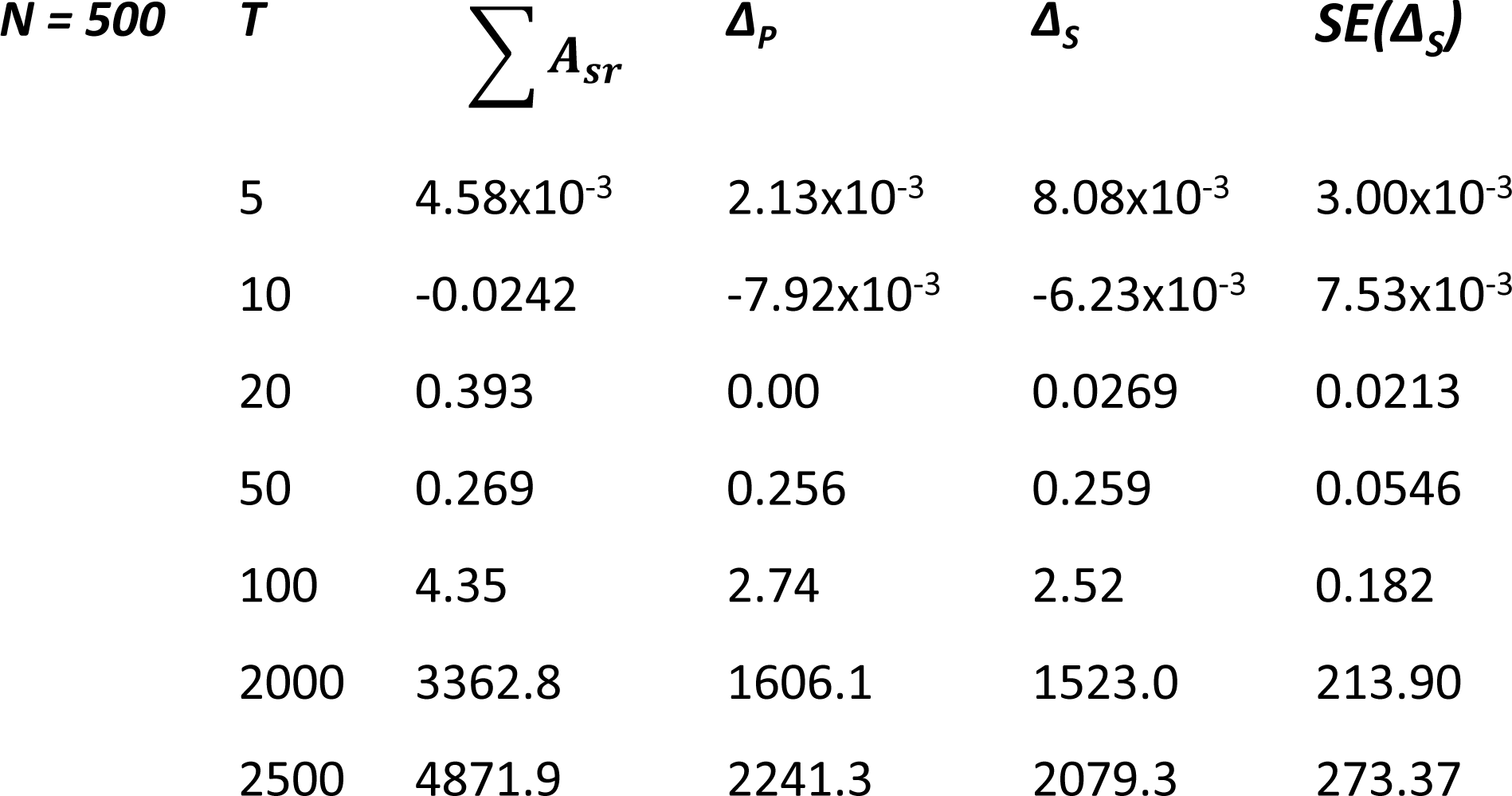
These tables show the number of segregating sites *S*_*n*_ in a sample of *n* = 20, the mean pairwise genetic distances 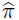*_HCS_*, 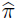*_LCS_* (for high and low coverage sequencing, respectively), and the variances in pairwise genetic distance for HCS vs. LCS. The latter are used to compute Δ*_S_* in Table 1. Table 2a shows these summary statistics for *N* = 200, Table 2b for *N* = 500.

Over *T ≪ N* generations, there are very few (~ 100) polymorphic sites, all of which have low variant allele frequencies. The mean and variance of genetic distances are of the order ~ 1, ~ 0.1, respectively. For *T ~* 4*N*, allele frequencies and genetic distances are near the equilibrium values, e.g. the estimated pairwise genetic distance 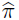 converges to the Tajima estimator for haploids *θ* = 2*Nu*, which is 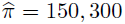 for *N* = 200, 500, respectively. Table 2 shows the sum of weighted LD values Σ*A*_*sr*_ = *Ā_sr_N*_2_. The mean values of *D*_*sr*_ are effectively 0 (~ 10*^−7^*10^−6^, either positive or negative, not shown in table) even for large values of *T* and *S*_*N*_. However, the skewed distribution results in a large postive weighted LD. Figures 2a-c show frequency distributions of allele frequencies, pairwise LD and weighted LD for a representative model population, specifically, for a single sample path where *N* = 500, *T* = 2500.

**Figure 2a-c.**
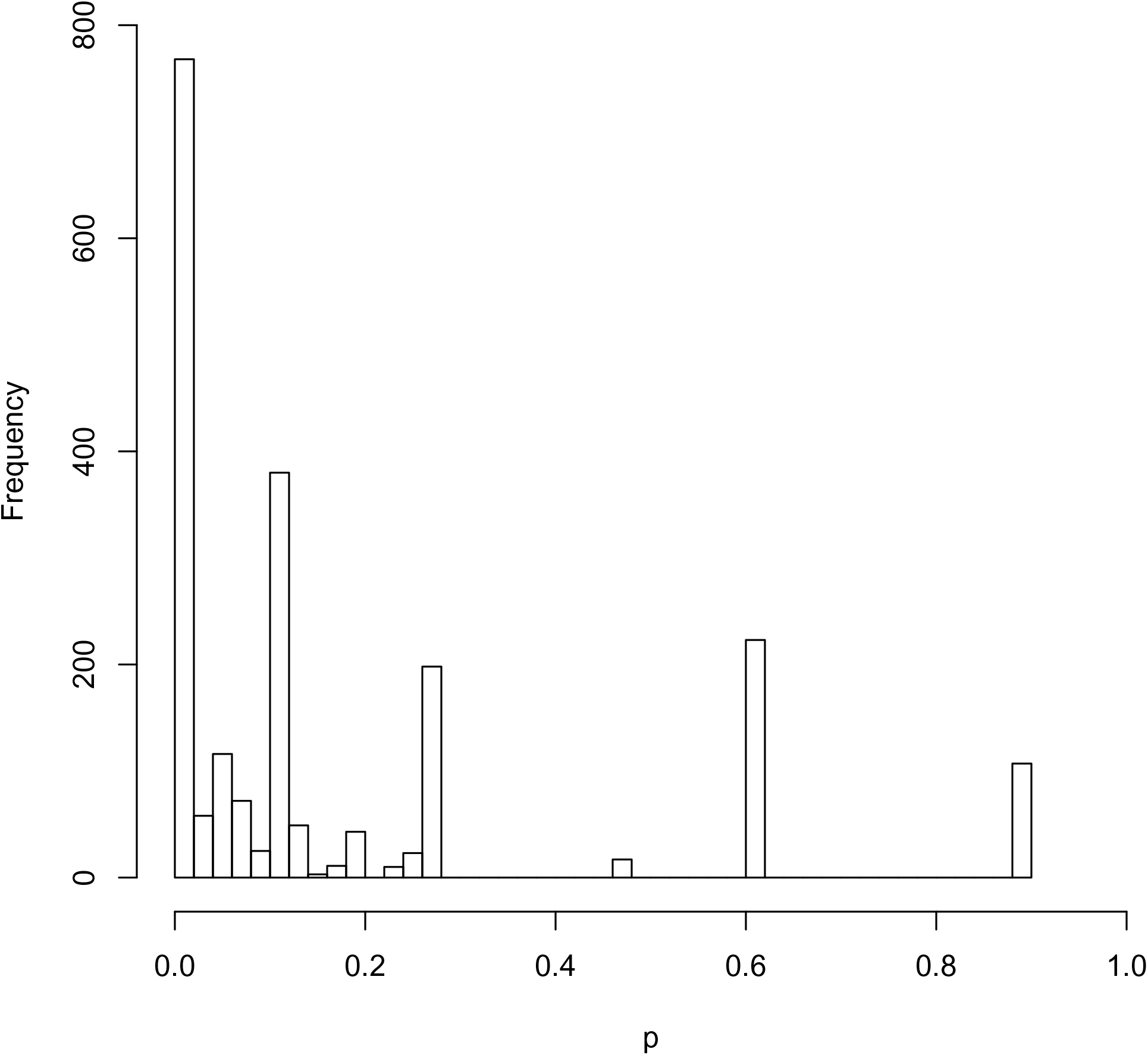
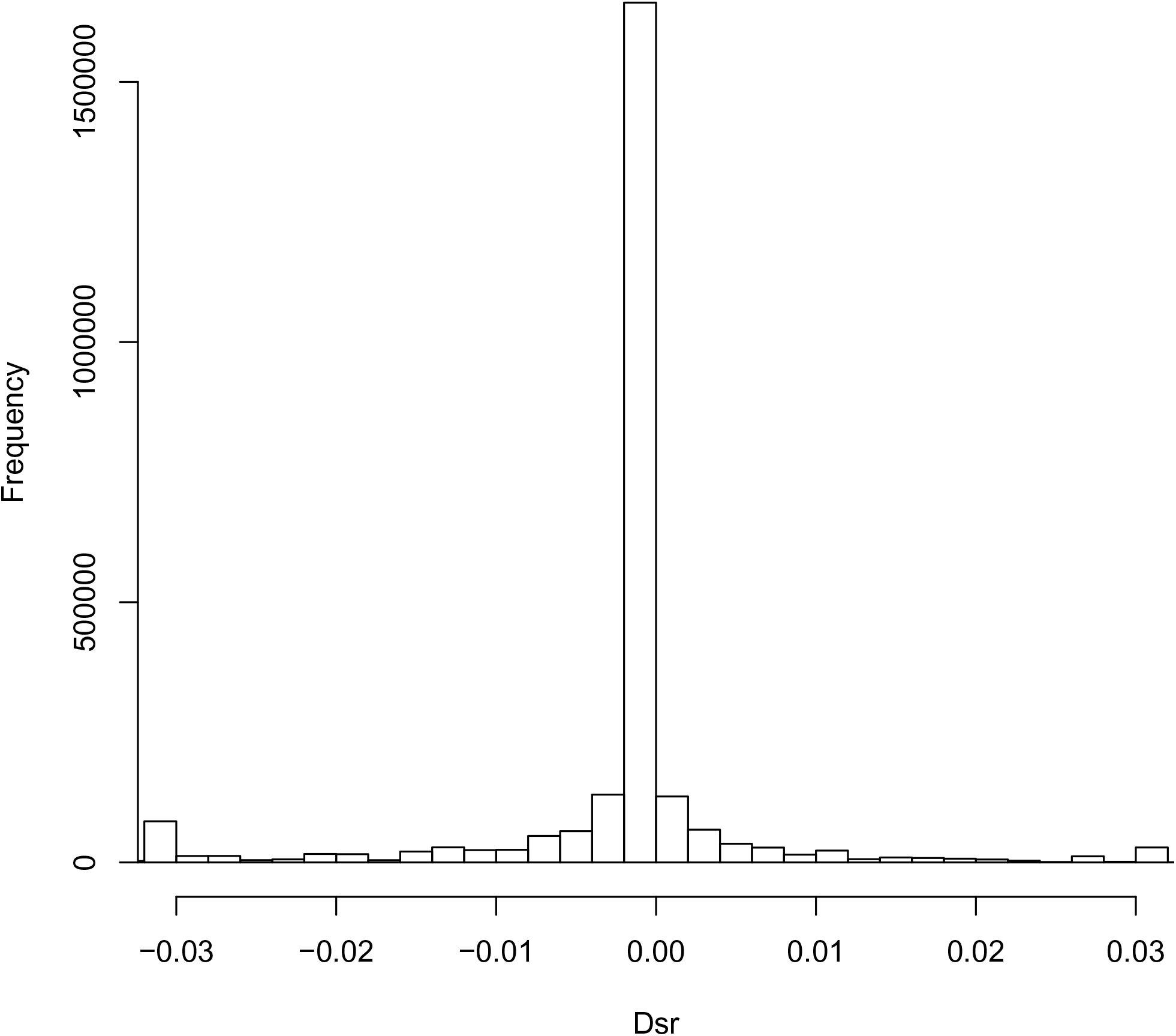
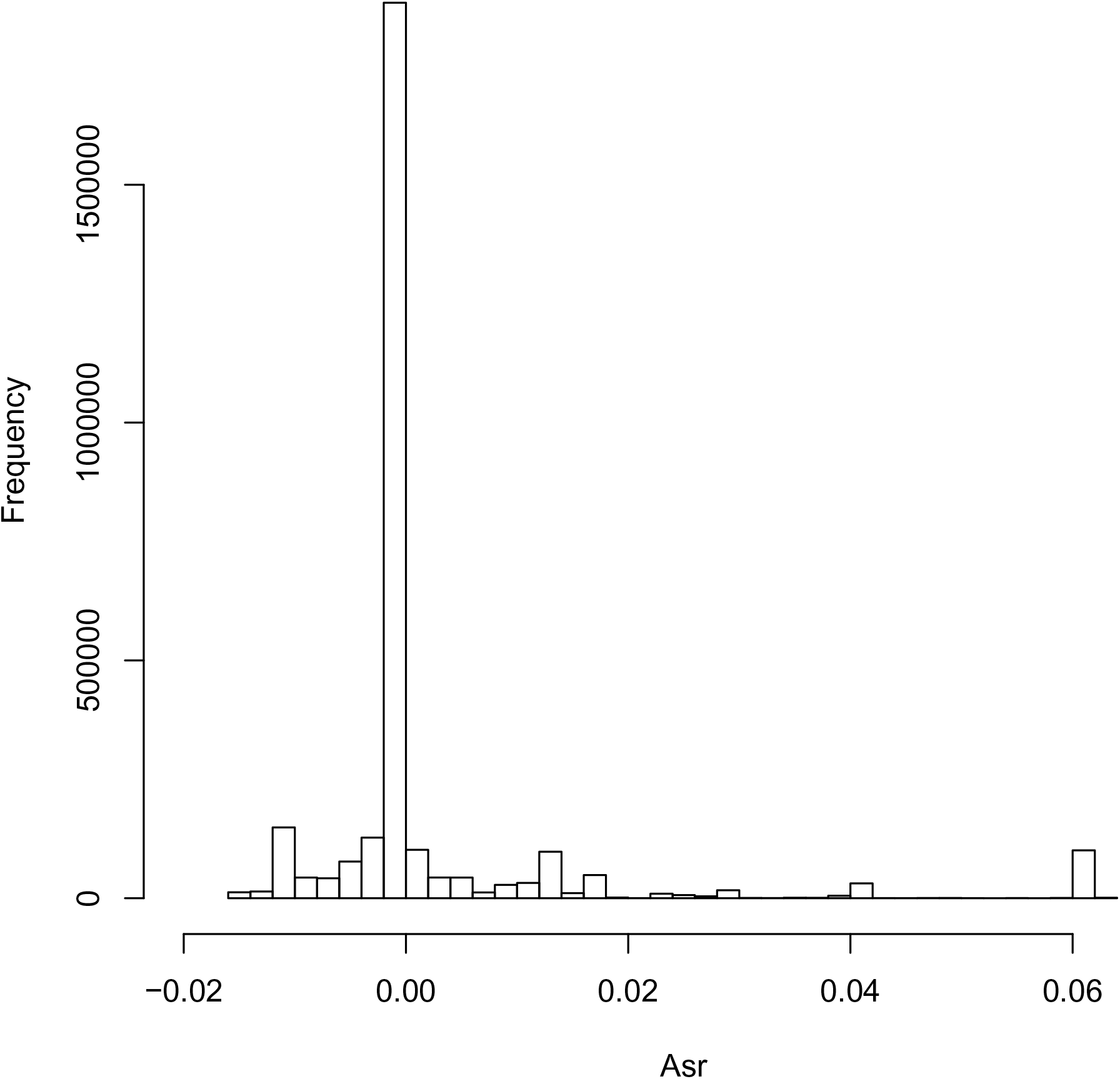
Figures 2a,b,c show, respectively, a representative distribution of variant allele frequencies *p*_*s*_, pairwise linkage disequilibria *D*_*sr*_, and weighted linkage disequilibrium *A*_*sr*_ for a simulated population with *N* = 500 haplotypes following *T* = 2500 generations of mutation and Fisher-Wright genetic drift. This is a single sample path (genealogy) rather than an averaged over all replicates, hence the outlier of high frequency mutations associated with a high frequency haplotype with a high density of mutations (Fig 2a). These in turn account for the positive skew (and positive mean value) of *A*_*sr*_ in a typical sample path (Fig 2c)

From Σ*A*_*sr*_, we compute the predicted difference between HCS and LCS variances Δ*_P_* using Eqn. (5). The predicted value is compared to the simulation estimate Δ*_S_* = *var_HCS_ − var_LCS_*. The consistency of observed and predicted values of Δ is confirmed by the fact that even the largest deviations are within less than two standard error 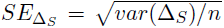 units with respect to the point estimate Δ*_S_*. The fit between analytical predictions and observed values improves for longer generation times as populations approach a mutation-drift equilibrium distribution of allele frequencies.

Except for populations with very few mutations where weighted LD values are ~ 0, we have Δ *>* 0 for most of the simulations. These results are consistent with LCS reducing the error in sample genetic distance except when variant alleles are rare and at low frequency. This reduction of error through low coverage sampling of many genomes is strongest for near-equilibrium distributions of allele frequencies, for large numbers of segregating sites, and for small sample sizes (corresponding to low coverage depth with NGS). Δ scales approximately as ~ 1*/n* for large *n*; consequently, for sample numbers and coverage depths of the order ~ 100, ∆ will be smaller by nearly an order of magnitude relative to the values obtained for *n* = 20 (simulations were performed for *n*= 10, 50, the results are not shown due to qualitative similarity to the data in Tables 1–2).

The two observed cases with *Ā*_*sr*_ < 0 are for *T* = 10, with a negative predicted value Δp for *N = 500* (though not for *N* = 200). Here, the Δ values are effectively zero within a standard error unit, so whether positive or negative values are observed is of purely formal interest (note that for even smaller time intervals *T* = 5 and even fewer polymorphic sites, both *Ā*_*sr*_ and Δ *>* 0, albeit very small). This suggests that at least under neutral evolution, *E(A*_*sr*_) < 0 occurs under restrictive conditions corresponding to very small absolute values of Δ and negligible reduction of error in estimating 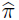 through either HCS or LCS, while for large numbers of segregating sites and increasing allele frequencies, there will be considerable increases in error when 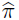 is estimated via high coverage sequencing.

For large *T*, the distribution of allele frequencies in the population approaches mutation-drift equilibrium, so that the mean *A*_*sr*_ should approximate *E(A_sr_)* computed from Eqns (7-9) using Mathematica 11.1. For *N* = 200, 500 and genomic mutation rate *u* = 0.3 (9 = 120, 300), the estimated values of *E(A*_*sr*_) = 1.11 × 10^−3^, 1.06 × 10^−3^, respectively, consistent in sign and magnitude with the mean values of *A*_*sr*_ for the simulated populations at *T* ~ 4N. For comparison, evaluating *Ā*_*sr*_ = *2*Σ*_s,r_ A_s,r_/S(S − 1)* for the number of polymorphic sites *S* (using the summed values of *A*_*sr*_ in Table 2, and the number of segregating sites *S* in Table 1), we obtain *A*_*sr*_ = 1.40 × 10^−3^ for *N* = 200, *T* = 1000 and Al_sr_ = 8.50 × 10^−4^ for *N* = 500, *T* = 2500.

The positivevaluesof *E(A_sr_)* result fromthefactthateventhough *E(_k_) ~ θ/k* suggests that high frequency alleles that contribute to large positive LD are rare, the high variance in allele frequencies means that any given sample path will usually contain several haplotypes characterized by multiple variant alleles at very high frequency. This means that even though the expectation for a particular *η*_*k*_ will be low for *k ~ N*, most genealogies will be characterized by a high count for some individual large value(s) of *k*, as is seen in Figure 2a. This generates a positive skewed distribution of pairwise linkage disequilibria (consistent with the distributions of *D*_*sr*_ derived numerically for non-recombining loci in (Golding, 1984)), as is seen in Figs. 2b-c. This contributes large positive *A*_*sr*_ values for most individual genealogies in the simulations, so consequently, *Ā*_*sr*_ > 0, Δ *>* 0. These results hold not only for populations at mutation-selection equilibria, but also for any populations where variant alleles have had time to accumulate to sufficiently high frequencies, hence the positive (albeit lower) values of Δ observed for all but the smallest time intervals.

## 5 Analysis of cancer sequence data

In this section, we apply the results of our derivations to genomic data by computing Σ*A*_*rs*_ and Δ for haplotype frequencies estimated from a lung adenocarcinoma sequences. Unpublished data on variant frequencies was provided to the authors by K. Gulukota and Y. Ji, who obtained their data via whole-exome sequencing of 4 sections of a primary solid tumor taken from a lung cancer patient. DNA from the samples was extracted using Agilent SureSelect capture probes. The exome library was sequenced with paired-end 100 bp reads on the Illumina HiSeq 2000 platform. Reads were mapped onto the human genome HG19 using BWA (Li and Durbin, 2009), giving a post-mapping mean 60-70 fold coverage across sites. Variant calls were performed with GATK (McKenna et al., 2010). Through the matching of read ends, somatic mutations co-occurring within ~ 100 bp in single genomes were identified (Sengupta et al. 2015). These mutation pairs define two locus haplotypes that can be tallied without the need for phasing, giving estimates of haplotype frequencies (defined by two proximate polymorphic sites) directly from the read counts.

Because reproduction in tumor cells is asexual and ameiotic, estimates of *D*_*sr*_ and *A*_*sr*_ using a subset of nearly adjacent sites is as representative of other haplotype pairs as if they were located on different chromosomes or on distant loci. The adenocarcinoma data contain estimated frequencies of 69 two-locus haplotypes, and corresponding variant allele frequencies for a total of 138 sites. This provides sufficient data to estimate the LD and weighted LD, and consequently the expected error in estimates of high versus low coverage sequencing.

A naive application of Eqn. (5) to the distribution of mutation frequencies and LD values gives Δ ~ 0.1 for coverage depth *n* = 65, suggesting much lower error if 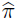 were estimated for the tumor via low coverage sequencing of pooled tumor cell genomes. However, several aspects of cancer genetics complicate this estimate. First, because cancer cells reproduce clonally, somatic mutations appear in heterozygous genotypes in the absence of mitotic recombination and gene conversion. A SNV frequency of *p* = 0.5 corresponds to fixation of a somatic mutation in a population of asexual diploids. Therefore, if we have heterozygous fixation at a single SNV site, a population consisting of 0/1 (reference and variant) genotypes, a mean genetic distance measure of 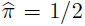 is meaningless because the population is homogeneous with respect to the 0/1 genotype, and variant allele frequencies must be rescaled to reflect this.

Figure 3 shows the distribution of mutant allele frequencies in Sample 1; note the high frequency of values near *p* = 0.5, the skew of the distribution is presumably the result of a low rate of detection of rare variants.

**Figure 3.**
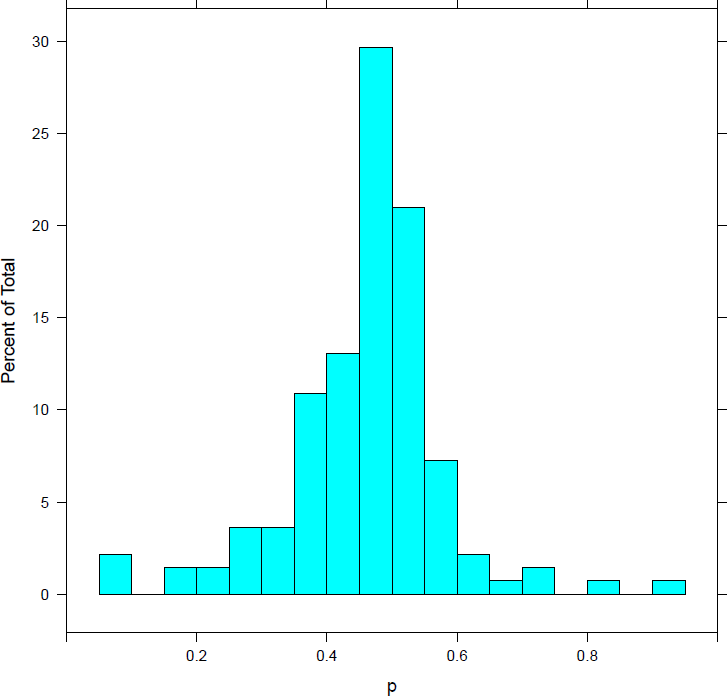
Distribution of allele frequencies *p* in the first lung adenocarcinoma sample, for *S*_*n*_ = 138 polymorphic sites. Values of *p* near 0.5 indicate heterozygous variant genotypes near fixation. Values *p >* 0.5 are a consequence of loss of heterozygosity via gene conversion during mitotic recombination, these are excluded from our analyses.

Williams et al. (2016a, b) (see also Ling et al. 2015) address the issue heterozygous genotype fixation by only considering polymorphic, segregating sites when comparing allele frequency spectra to neutral models, to the exclusion of sites that are ≥ 0.5 within a margin of sampling error; which also excludes sites whose frequencies *p >* 0.5 due to loss of heterozygosity. For the truncated range of allele frequencies *p* = [0, 0.5], the frequencies are rescaled to reflect heterozygosity, which for diploids means mapping *p^′^* = 2*p*, or more generally, *p^′^* = *p/f*_*c*_ where *f*_*c*_ is the cutoff for the inference of fixation. With this mapping, 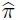 for a sample where all genotypes at a variant site are 0/1 is 0.

Assuming diploidy at all of the genotyped SNV sites and defining fixation as *p* = 0.5, we find that for sample size *n* = 65, the binomial probability of observing fewer than *x* = 26 mutant alleles is *Bin*(*x* ≤ 25*|n* = 65, *p* = 0.5) = 0.041, so we use *f*_*c*_ = 0.4 as as a cutoff defining polymorphic sites. By this criterion, and the rescaling *p^′^* = *p/f*_*c*_, there are only between 6 (sample4) and 10 (sample 3) adjacent segregating sites, and consequently between 3 and 5 haplotypes defined by such a pair out of the original 69. The LD and Δ values for this subset of haplotypes are summarized in Table 3. The differences in variances Δ remain positive, consistent with sample variance being lower with pooling. Δ is small (0.034 ≤ Δ ≥ 0.070), implying that in practice the estimation errors for 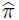 would be negligibly different between high and low coverage sequencing for this data set. However, the small Δ are partly a result of the small number of segregating sites (i.e. 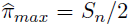) while 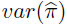 estimated by individual cell sampling may be expected to increase for more segregating sites, as was the case in the simulation data for larger time intervals.

**Table 3.**
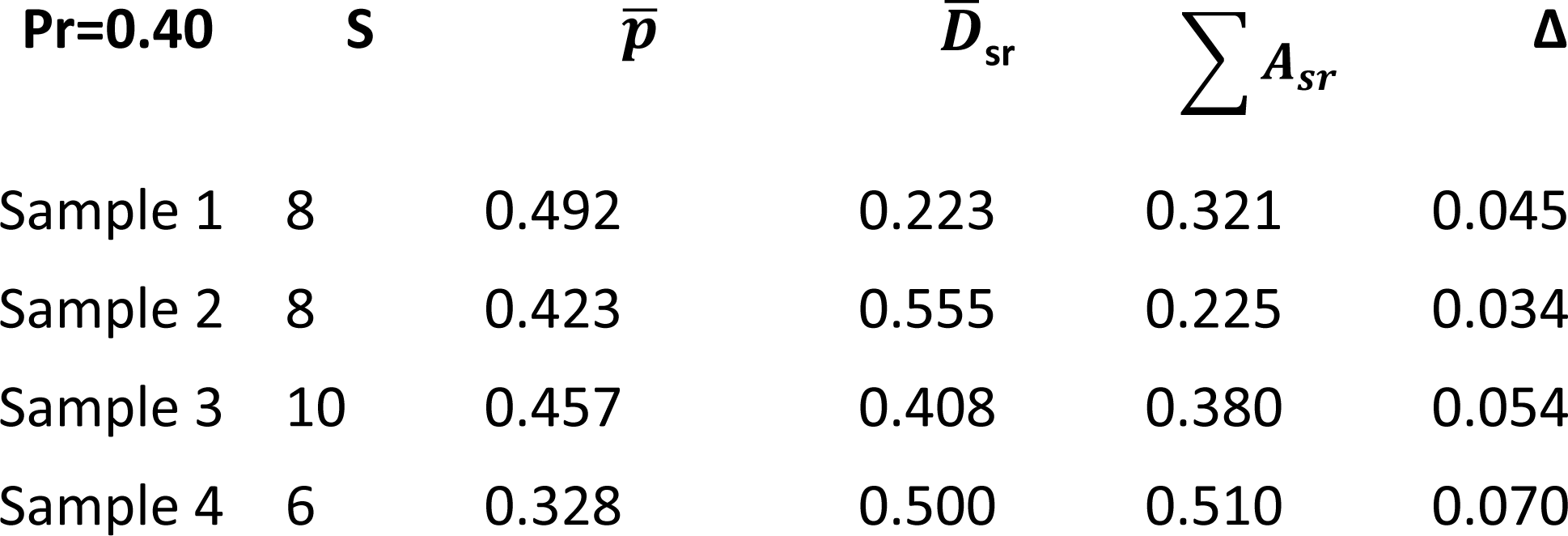
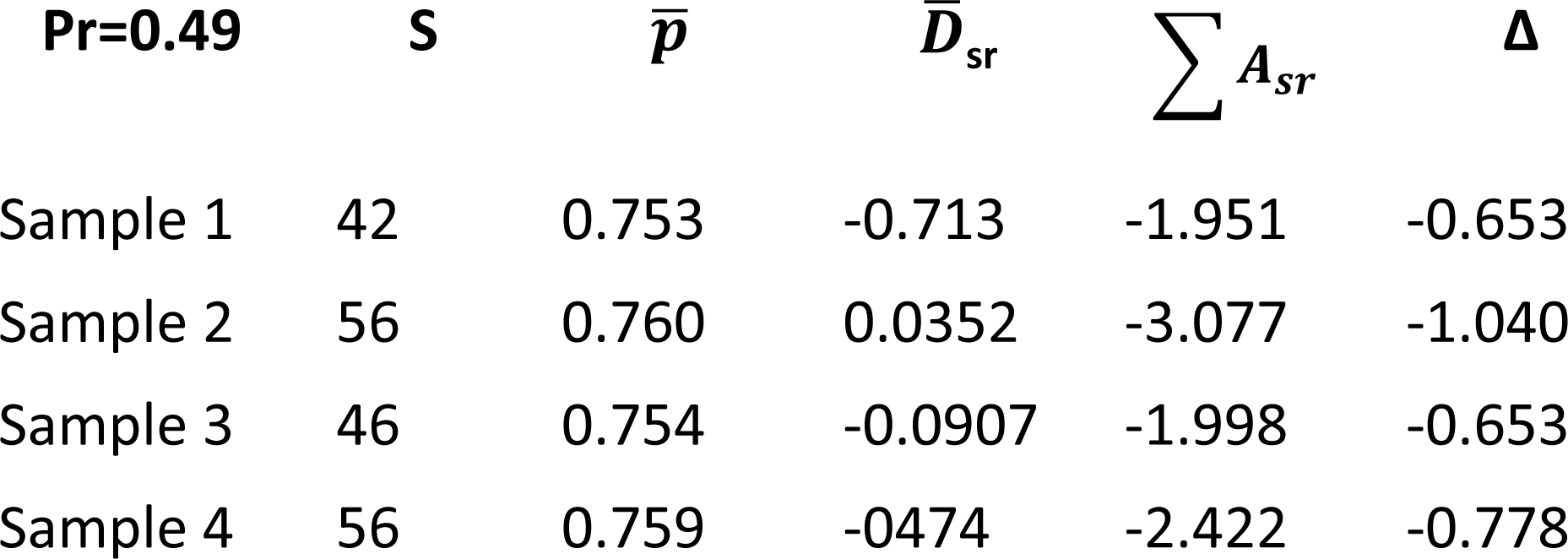
This table summarizes estimates of Δ from haplotype and allele frequencies in the lung adenocarcinoma sequence data, where haplotype frequencies for sites on individual long reads are known. Note that *Ā_sr_ >* 0 and Δ *>* 0 for all 4 samples, indicating that the error in pairwise genetic distance estimates for this data set are greater under HCS than under LCS, albeit weakly given the small number of unique haplotypes. Δ is computed from the mean read depth *n* = 65 for two cutoff values defining polymorphic sites. The upper panel shows the values for a cutoff of *f*_*c*_ = 0.40, selected based on a binomial probability. We use *p^′^* = *p/f*_*c*_, rescaled with respect to the diploid cutoff value. The lower panel shows the same for *f*_*c*_ = 0.49, selected arbitrarily close to *p* = 0.5 to show the sensitivity of Δ to the cutoff. The *f*_*c*_ = 0.40 calculations are based on 6-10 remaining polymorphic sites, the *f*_*c*_ = 0.49 on 42-56 sites, depending on the sample.

The values of Δ are also sensitive to the choice of truncation, as many of the SNVs occur in genotypes that are close to fixation in the tumor. For example, if we use *f*_*c*_ = 0.49, *x* = 32 as a cutoff to define segregating sites rather than *f*_*c*_ = 0.40, we obtain *Ā_sr_ <* 0 and Δ < 0 (of the order ~ 0.1). The sign reversal results from some lower frequency SNVs uniquely co-occuring in genomes with other SNVs that are close to fixation. The remaining allele and haplotype distributions contribute negative linkage disequilibria between the high frequency SNVs at one locus and high frequency reference alleles at the other site. The greater absolute value of Δ is a consequence of the fact that with a cutoff of *f*_*c*_ = 0.49, there are now 21-28 haplotypes (and 42-56 segregating sites) rather than the 6-10 for the *f*_*c*_ = 0.40 cutoff. The negative weighted LDs and Δ with this cutoff are shown in the second panel Table 3b, illustrating that for some samples, the variances in 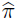 may actually be slightly higher with low coverage sequencing.

## 6 Discussion

The reduction of error in estimated genetic distance through low coverage sequencing reflects the loss of information due to non-independence across sites through linkage disequilibria. If the most frequent alleles at the majority of sites are in positive LD, any error in the estimated frequency and heterozygosity at one site covaries with the error at the other sites with individual sampling. In contrast, with LCS, each site provides independent information, so that the error across sites is uncorrelated. For *S*_*n*_ segregating sites in a sample of *n* and a variance in estimated distance per site *σ*^2^, with independent sampling the error across sites will approach *σ*^2^*/S*_*n*_. In the extreme case where allele frequencies across sites are nearly identical (complete linkage), the sample variance is *σ*^2^ independent of the number of sites. Another way to think of this is to consider the information gain that comes from sampling different loci from different subclasses of individuals in a sample under LCS, so that each polymorphic site has its own sample genealogy. This is analogous to the results of Pluzhnikov and Donnelly (1996), who found that in the presence of recombination, the optimal sequence length per genome for estimating allele frequencies and θ was sufficiently high to provide a large sample size while sufficiently short to provide low coverage per genome when recombination rates are low. With greater recombination rates, there is no information gain through low coverage because all but the closest loci have their own coalescent genealogy.

Conversely and by symmetry, a negative association of major allele frequencies across pairs of sites means that an error in estimated distance at one site will on average be compensated by an error in the opposite direction at another site, leading to reduction in variance under high coverage sequencing of individual genomes (analogous to improved estimation of the mean by sampling positive and negative extremes of a distribution). Both heuristic considerations and simulation results suggest that such a scenario is unlikely except for distributions of allele frequencies that give very small error values regardless, at least under neutral evolution. This was further confirmed by numerical calculation of the expected distribution of pairwise linkage disequilibria in an infinite sites model for clonal, ameiotic organisms.

Our results, like those of Ferretti et al. (2014) suggest that low coverage sequencing over pooled samples should be used to estimate the genetic distance (and consequently, population mutation rate parameter θ and the Tajima D statistic) under most conditions if reduction of estimation error is the sole criterion. However, there are several caveats to this conclusion, some theoretical, others practical. For example, we know that when most pairwise LD are approximately 0, the difference Δ between HCS and LCS estimates will be very small. A number of recent studies have shown that LD are generally among sites that are not physically linked in the genomes of sexually reproducing model organisms, including *Drosophila* (Andolfatto and Przeworski, 2000) and humans (Peterson et al., 1995; Reich et al., 2001). This suggests that any error introduced by sampling alleles from genomes individually with high coverage rather than pooled low coverage may be negligible for non-clonal genomes.

In contrast, in clonal organisms, or for regions of genome under very low recombination in sexually reproducing organisms, LD values will be high. Depending on the distribution of allele frequencies, Δ will be large when evaluated over many polymorphic sites. In the cases of cancer and microbial genomics, the standard NGS approach to sequencing reads from large numbers of cells at low coverage suggests an improved estimation of 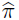 (and consequently, Δ and *N*_*e*_) relative to what would be obtained from more expensive single cell sequencing approaches. Furthermore, single-cell sequencing usually entails a much smaller sample size *n* than the coverage depths of 100-1000 that are standard for pooled sequencing. Moreover,Δ is defined on the assumption of the same effective sample size *n* for both LCS and HCS, when in fact LCS is associated with pooling and high read depth *n*, as is often the case, then this is often sufficient to reverse the sign of 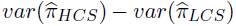 even in the rare cases when Δ *<* 0 for HCS sample size equal to LCS read depth *n*. This is because LCS combined with pooling increases the sample size per segregating site.

Finally, we remark that this study was to a large part motivated by efforts to apply the methods and theory of population genetics to cancer biology, where individual cell sequencing at high coverage versus pooled sampling with low coverage are often presented as alternative approaches. The case study computing Δ from lung cancer data in the previous section was used as proof of principle. A more accurate and refined analysis would have to take into consideration a number of potentially confounding variables. These include polyploidy and aneuploidy (so that with ploidy *X*, fixation corresponds to *p* = 1*/X*), as well as accounting for the loss of heterozygosity through mitotic recombination, reflected in frequencies *p >* 0.5. The sensitivity of 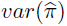 to the choice of cutoff *f*_*c*_ defining fixation for both the diploid and polyploid cases is of interest as an area for future research.

## 7 Acknowledgments

MS and JL were supported by the St. David‘s Foundation impact fund. YN and PM were supported by NIH grant 2R01CA132897-06A1. We thank Kalamakar Gulukota and Yuan Ji at NorthShore University HealthSystem for providing the lung cancer data used in this paper. We also thank two anonymous reviewers for their helpful critique and comments, as well as the following individuals for advice and assistance: Mark Kirkpatrick, Jon Wilkins, Matthew Cowperthwaite, Habil Zare, Mikhail Matz, and Andrea Sottoriva.

## 9 Appendix A1: Ordered Pairs of Pairs

Recall the definitions *µ* = *E*(*ϕ*_*ij*_), *σ*^2^ = *var*(*ϕ*_*ij*_) and *κ* = *E*(*ϕ_ij_, ϕ_jk_*).

**Lemma 1.** *Let µ* = *E*(*ϕ*_*ij*_) *where the expectation is over pairs x_i_ ~ p*(*x*) *and x_j_ ~ p(x), independently. Let* 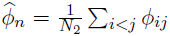, *denote a sample estimate for µ, averaging over all pairs* (*i, j*) *of samples. Then* 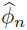 *is unbiased*, 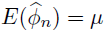, *and*

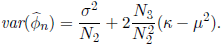

*Proof.* Unbiasedness is straightforward:

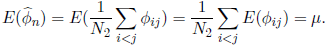

For the variance, note that

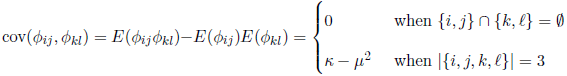

Then

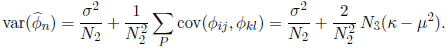

*Proof of Eqn. (3)*. Let 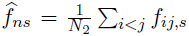. From the statistical independence among sites *s* located on different reads under LCS, it follows that for a sample of *n*,

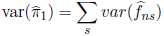

with

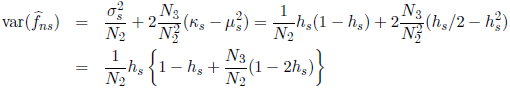

where the first equality is due to Eqn. (2).

## 10 Appendix A2: Poisson Distribution of Read Depth

In practice, the read depth with LCS is not constant and varies considerably across sites. Assuming the read depth follows a Poisson distribution with mean *n* equal to the number of haplotypes in HCS. Computing the Taylor expansion of the variance of genetic distance estimator, we find the difference between fixed and Poisson read depth is 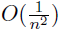.

Let *n*_*s*_ denote the read depth at locus *s* which follows a Poisson distribution *n_s_ ~ P oi*(*n*) with mean *n*. Let

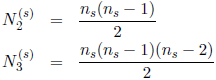

Let 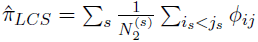 and ***n*** = (*n*_*s*_)*_s_*. The variance of 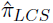 conditioned on the sampling depths ***n*** is given by

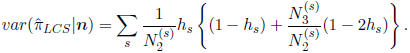

By the law of total variance, we have

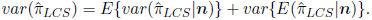

The second term is zero because the expectation is independent of ***n***,

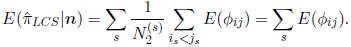

The first term is non-trivial. Let 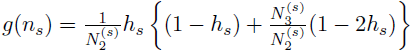 and expand *E*(*g*(*n*_*s*_)) at *n*,

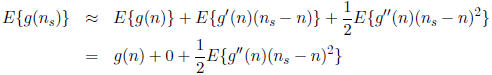

with

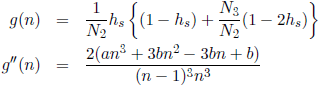

where *a* = 2*h*_*s*_(1 *−* 2*h*_*s*_) and *b* = *−*2*h*^2^*_s_* + 6*h*_*s*_. So

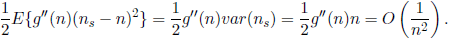

Therefore,

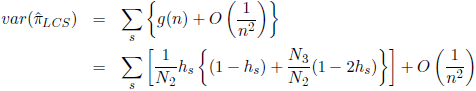

where the first term of the last equality is the variance of 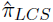 given the read depth is fixed at *n* across all loci. This result holds for any sampling distribution of read depths where the variance is of the order *~ n*.

The very small 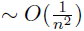 contribution of non-constant read depth to the error in estimated 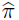 is confirmed using simulation results. Compare the variances and other parameter estimates observed in Table 2 for *N* = 200, *n* = 20 where the read depths are constant to those in Table A2, where the read depth is a Poisson random variable with parameter *n*.

**Table A2.** This table summarizes the same parameter estimates as in Table 2a (for *N* = 200, *T* = 1000), however, the LCS read depth is now a Poisson random variable with mean *n* = 20 rather than a constant. Note that the simulation values for the means and variances of 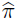 and of *A_sr_,* Δ are largely unchanged due to the negligible contribution of variance in read depth to the error in parameter estimation.

| *N=200* | *T* | $\sum A_{sr}$ | $\Delta_p$ | $\Delta_s$ | $SE(\Delta_s)$ |
| --- | --- | --- | --- | --- | --- |
|  | 5 | 2.82x10<sup>-3</sup> | 5.76 x 10<sup>-4</sup> | 3.71x10<sup>-4</sup> | 0.071 |
|  | 10 | 0.0541 | 0.0250 | 0.0242 | 0.014 |
|  | 20 | 0.0181 | 0.0696 | 0.0704 | 0.034 |
|  | 50 | 1.232 | 0.792 | 0.620 | 0.080 |
|  | 800 | 754.33 | 341.14 | 296.18 | 42.59 |
|  | 1000 | 827.67 | 373.24 | 337.70 | 57.88 |

